# Sugar chain structure of apolipoprotein B-100 and its role in oxidation

**DOI:** 10.1101/2022.05.31.494124

**Authors:** Masahiko Okada

## Abstract

The aim of this research was to determine the structure of oligosaccharide antennae located on the surface of apoB-100, and to examine their roles in the oxidation process and in the signal transduction of endothelial cells. The profiles of oligosaccharides on apoB-100 were determined by enzymatic digestions as follows. First, N-glycanase was used to release a mixture of oligomannose and complex types of oligosaccharides. Second, endoglycosidase H was used to release high-mannose and hybrid types. Third, O-glycosidase DS was used to release O-linked oligosaccharides. The released oligosaccharides were then labeled and quantified by electrophoresis. In vitro apoB-100 oxidation was mimicked by adding transition copper ions. For the signal transduction study, I examined the expression of adhesion molecules on cultured human coronary artery endothelial cells by adding LDL in which the oligosaccharide sequences were enzymatically modified. The sugar chain structures on the surface part of apoB-100 were composed predominantly of N-linked oligosaccharides, i.e., two forms of complex type and five forms of high-mannose type. The digestion of sugar chains by exoglycosidases and endoglycosidases did not result in any changes in the susceptibility of LDL to oxidation. Also, LDL without monosaccharides such as sialic acid, galactose, and N-acetylglucosamine did not induce any significant effect on the expression of ICAM-1, VCAM-1, or ELAM-1. I found that the sugar chains did not play any significant roles in the oxidative processing of LDL and also in the expression of ICAM-1, VCAM-1, or ELAM-1.

---

Increasing evidence has shown that sugar chain structures are often important as recognition determinants in receptor-ligand or cell-cell interactions, for example, in the modulation of immunogenicity and protein folding, and in the regulation of protein bioactivity. In general, sugar chains are composed of monosaccharides, such as sialic acids, galactose and mannose, and exist as a combination of oligosaccharide antennae with various structures.^1^ Apolipoprotein B-100 (apoB-100) of low- density lipoprotein (LDL) is a glycoprotein that is composed of such oligosaccharides.^2^

The oxidation of apoB-100 is widely believed to be an important event in the pathogenesis of atherosclerosis. LDL with oxidized apoB-100 is endocytosed in an uncontrolled manner by macrophages, resulting in the generation of cholesterol-laden foam cells, which characterize atherosclerotic lesions.^3, 4^ Sugar chain structures may play a role in the oxidation process of the glycoproteins or in the signal transduction of endothelial cells.

The aim of this research was to determine the structure of oligosaccharide antennae located on the surface of apoB-100, and to examine their roles in the oxidation process and in the signal transduction of endothelial cells. In vitro apoB-100 oxidation was mimicked in a cell-free system by adding transition copper ions, as prooxidants.^5–7^ For the signal transduction study, I examined the expression of adhesion molecules on cultured human coronary artery endothelial cells by adding LDL in which the oligo-saccharide sequences were enzymatically modified.

## Materials and methods

### Samples

Plasma samples (3.4 mmol/L K_2_-ethylene-diamine tetraacetic acid (K_2_ EDTA)- anti-coagulated venous blood) were obtained from subjects with no obvious arteriosclerotic diseases and stored at 4 C before experiments. The study protocol was approved by the Committee on Human Research, Niigata University School of Medicine, Japan, and written informed consent was obtained from all subjects.

### LDL Preparation and Oxidation

LDL (1.019 < *d* < 1.063 μg/ml) was prepared with an ultracentrifugation method as follows. One ml of a solution of sodium bromide (1.025 μg/ml) supplemented with 0.01 % (w/ w) Na N_3_, 0.3 mmol/L Na_2_ EDTA, and 1 mmol/L benzamide hydrochloride were added gently to 2 m L plasama, and the mixture was ultracentrifuged at 400, 000 × *g* (at the bottom of tube) at 4 °C for 5 h. Two ml of the bottom fraction were then transferred to another tube, and the density was increased to 1.063 with a Na Br solution. After another centrifugation at 400,000 × *g* at 4 ° C for 5 h, the top fraction was collected.

Prior to oxidation, each LDL fraction was applied to a Sephadex G-25 column (PD-10; Pharmacia Biotech, Uppsala, Sweden) for exchanging buffer with phosphate buffered saline (PBS). The protein content of the LDL sample was determined by a protein assay kit (Bio-Rad Laboratories, Inc., Tokyo, Japan) based on the Bradford method using bovine serum albumin as a standard. Then, 4 mol/L Cu SO_4_ (final solution) was added to each aliquot, incubated for 5 h, and the oxidation was stopped by separating the copper ions from LDL using a PD-10 column.

The electrophoretic mobilities of both native and oxidized LDL were examined for each sample to confirm successful oxidation. Electrophoresis was carried out on 1.0 % agarose gel films (TITAN GEL Lipoprotein, Helena Laboratories, Saitama, Japan) in a 0.08 mol/L barbital buffer at p H 8.6 and run at 90 V for 25 min, according to the manufacturer’ s instructions. Lipoproteins were visualized by staining with 0.04 (w/v) Fat Red 7 B in 95 % methanol. The protein amount in each applied sample was adjusted to 100 μg/mL (final concentration). The relative mobility of the LDL bands was automatically measured by distance in millimeters with an AE-6981 GXCP ATTO COMBO II imager (ATTO, Tokyo, Japan). The ratio of the distance of oxidized LDL to that of native LDL was obtained.

### Oligosaccharide Profiling and Sequencing

The profiles of oligosaccharides on apoB-100 were determined by enzymatic digestions as follows. First, N-glycanase was used to release a mixture of oligomannose and complex types of oligosaccharides. Second, endoglycosidase H (Endo H) was used to release high-mannose and hybrid types. Third, O-glycosidase DS was used to release O-linked oligosaccharides. The released oligosaccharides were then labeled with a fluorophore 8 - aminonaphthalene-1, 3, 6 - trisulfonic acid (ProZyme, Inc., San Leandro, CA) at the reducing termini by reductive amination. The fluorescent labeled oligosaccharides were separated and quantified by electrophoresis using an OLIGO Profiling Gel and a FACE imaging system (ProZyme, Inc.). The stoichiometry of labeling was such that only one molecule of fluorophore was attached to each molecule of oligosaccharide. The standard used in the analysis consisted of a mixture of glucose polymers ranging from a monomer to a polymer of 16 molecules (glucose_1_ - glucose_16_) (ProZyme, Inc.). Oligosaccharide bands in each sample were quantified by comparing the intensity of the glucose_4_ band (containing 25 pmols), of which the intensity in the standard mixture was prepared as intentionally weaker than adjacent bands. Visualized individual oligosaccharide bands were carefully excised using a razor blade. The oligosac-charidesthus isolated were then sequenced through a series of exoglyco sidase digestions and another electrophoresis. In detail, sialidase A (sialidase), (1-4) galactosidase (GALase), - N-acetyl-hexo-saminedase (HEXase), (1-2, 3, 4) mannose-dase (Jack bean) (MANase), and (1-6) mannosidase (recombinant from *Xantbo-monas manibotis*) (MANase) were sequentially added to each aliquot of the oligosaccharides. The degradation patterns were compared with the pre-determined migration patterns of oligosaccharides of known sequencesthat were provided by the manufacturer. The endoglycosidases and exoglycosidases were all purchased from ProZyme, Inc., except for O-glycosidase DS (Bio-Rad Laboratories, Inc., Tokyo, Japan). Three samples were analyzed separately. Modifications of the sugar chains of native apoB-100 were performed by adding each exoglycosidase sequentially. The amount of each enzyme needed to catalyze the oligosaccharides was determined experimentally.

### Cell Culture

Endothelial cells of a human coronary artery (the 3 rd passage) and endothelial growth medium CC-3125 (Cambrex Bio Science Walkersville, Walkersville, MD) were purchased from Sanko Junyaku (Tokyo, Japan). Endothelial cells were proliferated in 21 - cm^2^ or in 55 - cm^2^ culture dishes until reaching subconfluent monolayers (about 80 %) in the growth medium at a 0.15 m M calcium concentration supplemented with 5 % fetal bovine serum, 12 μg/ml human recombinant epidermal growth factor, 1 μg/ml hydrocortisone, 12 μg/ml bovine brain extract, 25 μg/ml gentamycin, and 25 μg/ml amphotericin B. For m RNA analysis, the cells in a 21 cm^2^ culture dish were washed by HEPES buffer, harvested by trypsinization (0.025 % trypsin and 0.01 % EDTA), received subsequent trypsin inhibitor treatment, centrifuged, and resuspended in a PBS.

Total RNA was obtained by using an RNA Isolation kit (Roche Diagnostics, Tokyo, Japan), which uses a glass-fiber fleece method, according to the manufacturer’ s instructions. At the time of the experiments, the expected number of cells in each dish was 3 × 10^6^.For protein analysis, cells in a 55 - cm^2^ dish were washed in PBS and then lysed for 15 min at 4 C in 500 L of Cel Lytic M buffer (Sigma, St. Louis, MO). After centrifugation, the supernatant was collected, condensed, and protein contents were assayed.

### Stimulation of Endothelial Cells

To compare the response of endothelial cells to a standard stimulant, I used 10 ng/ m L recombinant human interleukin 1 (IL-1α; R&D Systems, Minneapolis, MN). All incubations were done at 37 °C, p H 7.4, in a humidified atmosphere containing 5 % CO_2_ and 95 % air.

### Real-time Polymerase Chain Reaction

In order to quantify the differences in each m RNA expression, I analyzed samples by real-time reverse transcriptase polymerase chain reaction (real-time RT-PCR) using a LightCycler 1.5 (Roche Diagnostics). Measurements form RNA of vascular cell adhesion molecule-1 (VCAM-1) and β-actin were conducted using the following conditions: 10 min at 55 C and 30 s at 95 C for reverse transcription; 45 cycles of 1 s at 95 °C and 30 s at 60 °C for amplification; and 1 s at 95 °C, 10 s at 65 °C, and 1 s at 95 °C for melting curve analysis. Measurements for the m RNA of intercellular adhesion molecule-1 (ICAM-1) and endothelial leucocyte adhesion molecule-1 (ELAM-1) were conducted using the following conditions: 5 min at 42 °C and 10 s at 95 °C for reverse transcription; 40 cycles of 5 s at 95 °C, 20 s at 60 °C (ICAM-1) or 62 °C (ELAM-1) for amplification; and 1 s at 95 °C, 15 s at 65 °C, and 1 s at 95 °C for melting curve analysis. Sequences of the primers used are listed in Table 1 of the supplementary appendix.

The relative quantification of the expressions of the target genes was measured using β-actin m RNA as an internal control. The results were quantified by calculating fold increase as follows:

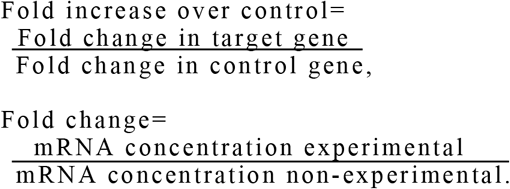

Every experiment was carried out in duplicate.

### Weatern Blot Analysis of Adhesion Molecules

Western blot analysis was performed using monoclonal antibodies against ICAM-1 (MBL, Nagoya, Japan) (concentration: 0.4 μg/ml), VCAM-1 (Santa Cruz Biotechnology, Inc., Santa Cruz, CA) (concentration: 0.4 μg/ml), ELAM-1 (Santa Cruz Biotechnology, Inc.) (concentration: 0.4 μg/ml) or β-actin (Abcam, Cambridge, MA) (concentration: 0.1 μg/ml). Two separate experiments were carried out.

### Statistical Analyses

Statistical significance was determined by unpaired Student *t*-test with or without Bonferroni correction. Differences were considered significant at *P* < 0.05.All statistical analyses were conducted using SPSS software, version 11.0 J for Windows (SPSS Japan, Tokyo, Japan).

## Results

### Sequence of Oligosacharides on LDL

In the electrophoresis for oligosaccharide profiling, two bands were identified after N-glycanase treatment and five bands were identified after Endo H treatment, but no bands were identified after O-glycosidase DS treatment (Fig. 1).The results showed that the sugar chain consisted of two different forms of complex type, five different forms of high mannose and/ or hybrid types, and no O-linked oligosaccharides.

**Fig. 1.**
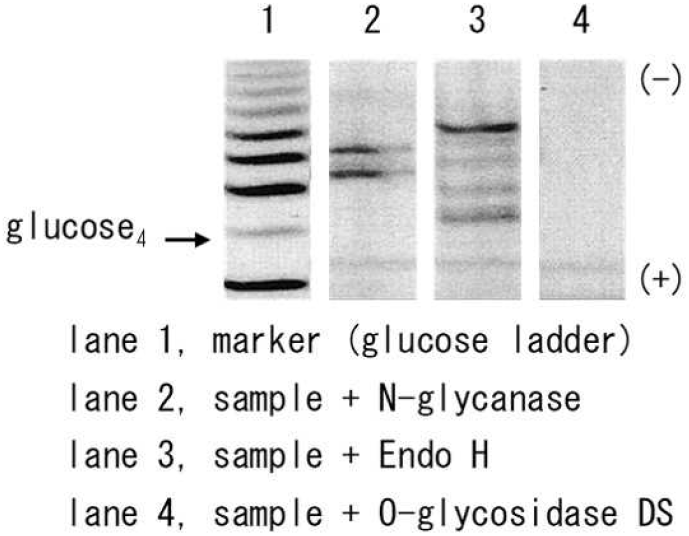
Oligosaccharide profiling. Lane 1 indicates glucose polymers ranging from a monomer to a polymer of 16 molecules (glucose_1_ - glucose_16_).Glucose_4_ contained 25 pmols of glucose polymer. Lane 2, Effect of N-glycanase treatment; Lane 3, Effect of Endo H treatment; Lane 4, Effect of O- glycosidase.

The upper band that was obtained by N-glycanse treatment (an N-linked oligo-saccharide) wasthen isolated and treated with sialisase. Migration values for oligosaccharides were defined in terms of their degrees of polymerization (DP). DP values were assigned to unknown oligosaccharides by comparison with the glucose ladder. Each oligomer in the glucose ladder corresponds to a specific DP value as follows: glucose_1_ = 1 DP, glucose_2_ = 2 DP, etc. After the treatment, the band migrated 1.44 DP upward, which demonstrated that a negative charge was removed (Fig. 2 A). Since a sialic acid residue is negatively charged and contributes an average of 1 DP unit shift, it could be determined that one of the first monosaccharides was a sialic acid. The lower band wasthen treated with a series of enzymes: sialidase, GALase, HEXase, and MANase (Fig. 2 B). I identified that the oligosacchardes were comprised of two sialic acids (NeuAc) and two galactose (Gal) residues. Since one antenna (the upper band) is composed of the same sequence plus one sialic acid residue, the number of antennae in N-linked oligosaccharides was identified as two. Two N-acetylglucosamine (GlcNAc) and two mannose (Man) residues were also identified. The core was not fucosylated.

**Fig. 2.**
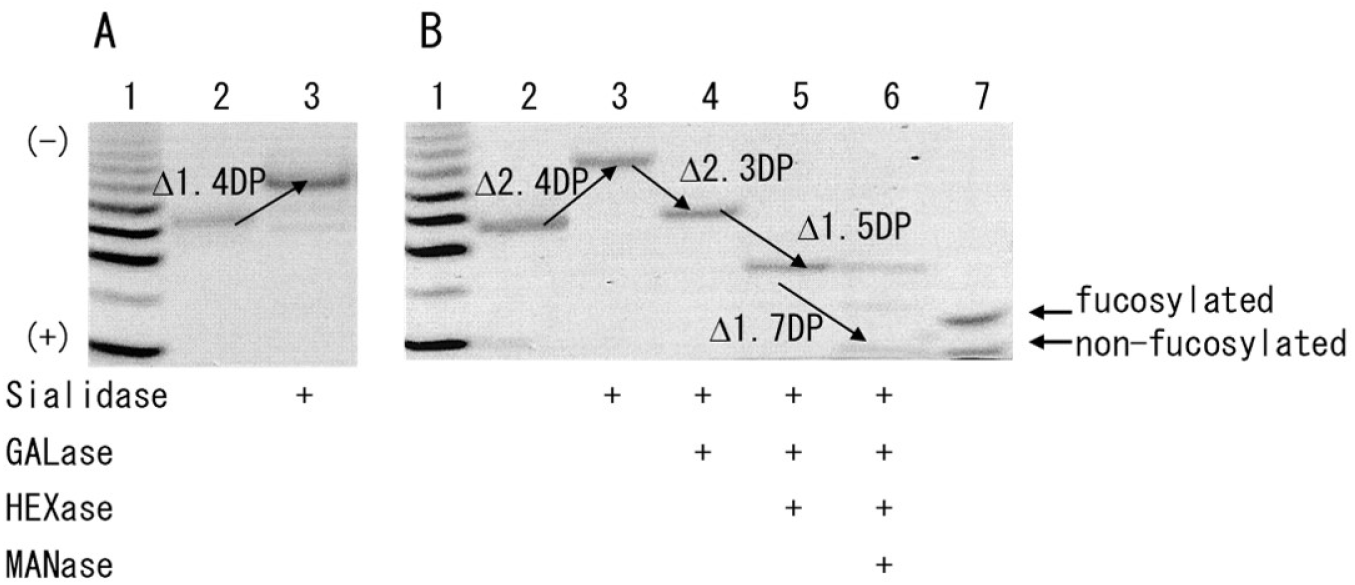
Oligosaccharide sequencing. A, Upper band of N-linked oligosaccharides (see Figure 1, lane 2) was treated with sialidase. B, Lower band was treated with a series of sialidase, GALase, HEXase, MANase (Jack bean), and MANase (Xantbomonas manibotis).

The estimated structures were

NueAcα2-6Galβ1-4GlcNAcβ1-2Manα1-6

(NueAcα2-3 Galβ1-4GlcNAcβ1-2Manα1-3)

Manβ1-4GlcNAcβ1-4GlcNAc and that without a NeuAc.

The five bands that were obtained by treatment with Endo H exhibited no diges-t ion until MANase was added. Therefore, these bands represented five different forms of high-mannose-type oligosaccharides and were not of hybrid types. I identified that these oligosac-charides contained between five and nine mannose residues attached to the chitobiose core. The estimated structures were

((Manα1-2)_0-4_) Manα1-6Manα1-6

(Manα1-3) Manβ1-4GlcNAcβ1-4GlcNAc.

Figure 3 shows the estimated structure of the sugar chains. There were no significant differences among samples obtained from three subjects.

**Fig. 3.**
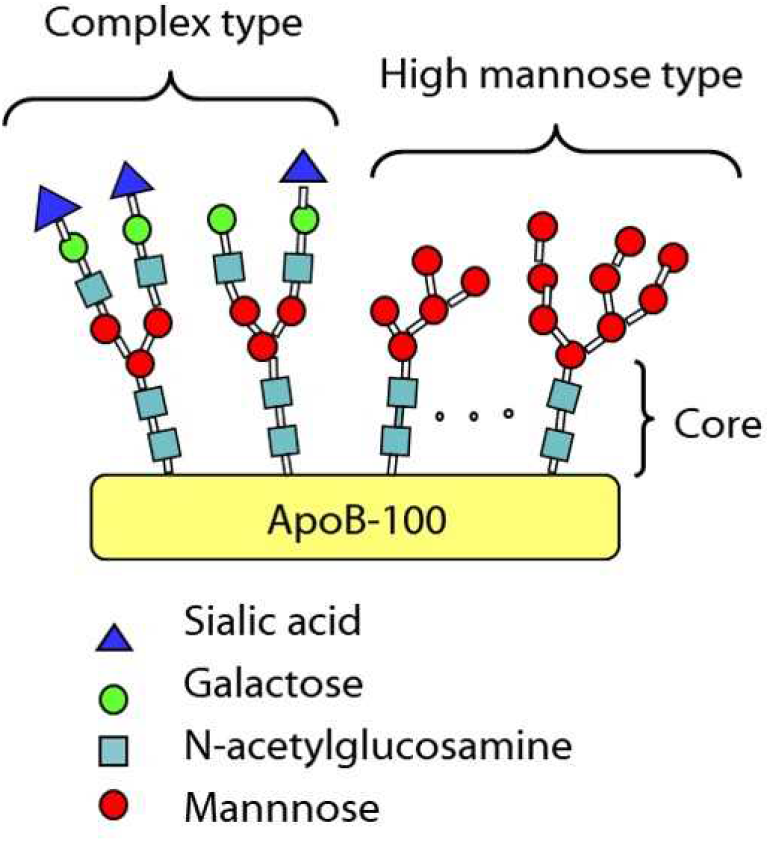
Proposed structure of sugar chains on apoB-100. Sugar chains are composed predominantly of two forms of complex type and five forms of high mannose type oligosaccha-rides.

### Oxidizability of Oligo-saccharide Modified LDL

Aliquots of the LDL sample were processed with a series of exogly-cosidases and incubi-ted for 5 h with 4 mol/L Cu SO_4_.There were no significant differences in the agarose gel electrophoretic mobilities, i.e., in the oxidizabilities (Fig. 4).

**Fig. 4.**
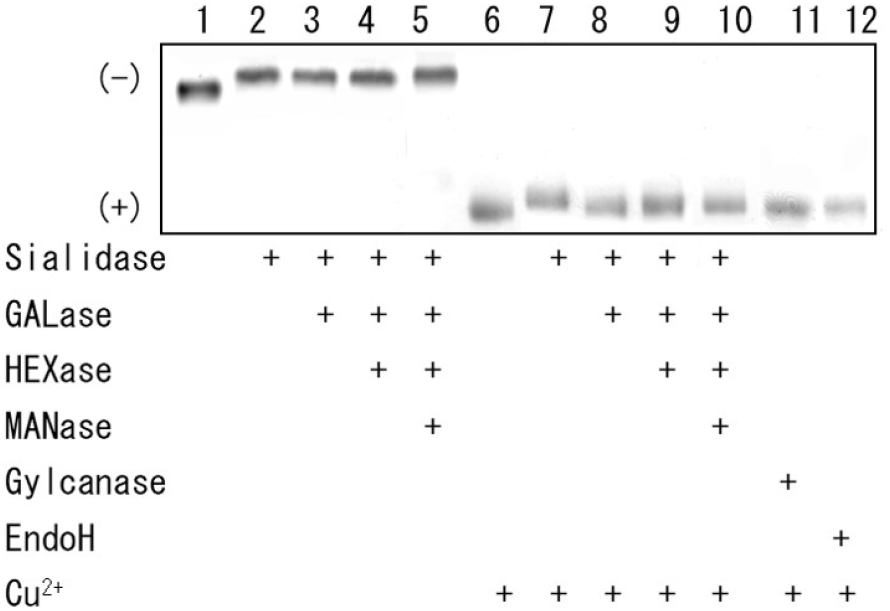
Oxidization of modified LDL. Aliquots of the LDL sample were processed with a series of sialidase, GALase, HEXase, MANase, N- glycanase, and Endo H, and incubated for 5 h with 4 mol/L Cu So_4_.Shown are electrophoretic mobilities on an agarose gel film.

### Expression of Adhesion Molecules

Adhesion molecule expressions were first examined under basal conditions and after a 24 - h incubation of cells with 10 ng/ml IL-1α in order to have positive controls for mRNA an d protein analyses. ICAM-1, VCAM-1, and ELAM-1 were constitutively expressed on the cells and super-induced by the cytokine treatments (Fig. 1 of the supplementary appendix). There was no significant change in the protein expression of β-actin between untreated and treated conditions. These results are compatible with previous studies.^8–13^

No significant increase was observed in the mRNA levels of ICAM-1, VCAM-1, or ELAM-1 by incubation with the LDLs that were treated sequentially with each exoglycosidase relative to the control (Fig. 5).

**Fig. 5.**
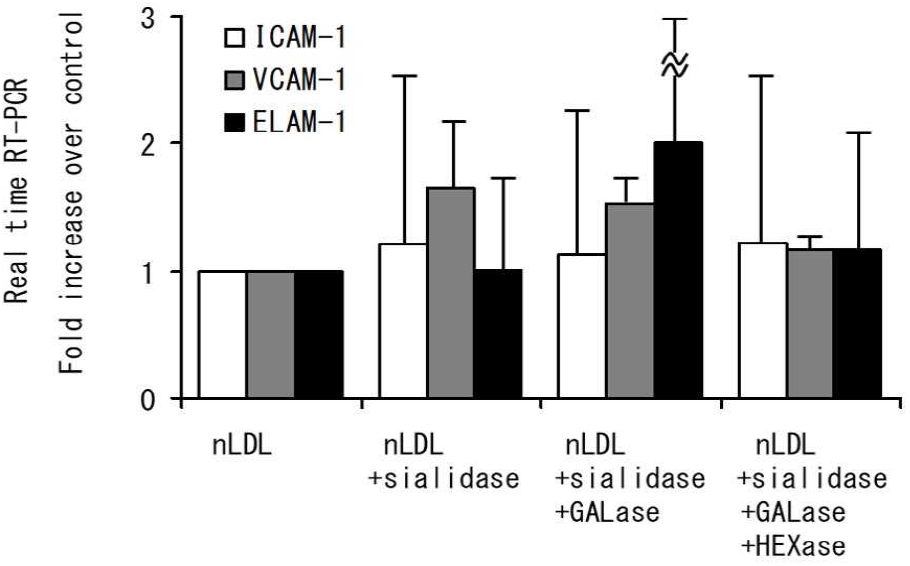
Effects of modified LDL on adhesion molecule expressions. No significant increase was observed in the mRNA levels of ICAM-1, VCAM- 1, or ELAM- 1 by incubation with the LDLs that were treated sequentially with each exoglycosidase relative to the control.

## Discussion

Sugar chains or oligosaccharides are initially added to a protein in the endoplasmic reticulum (ER) and are subsequently modified through enzymatic biosynthesis.^14^ The extent of branching or elongation of oligosaccharides differs from molecule to molecule. Hence, sugar chains do not have a uniform structure. The linkage between carbohydrate and protein is indicated by the locants N- and O-. The locant N-is used for the N-glycosyl linkage to asparagine. N-linked oligosaccharides are divided into two major classes: complex type and oligomannose type. Structures containing both classes are designated as a hybrid type. In the ER, a block of oligosaccharides is initially synthesized as a high-mannose type and maturesthrough various enzymatic modifications into a complex- or hybrid type. The complex type is the most mature among the N-linked oligosaccharides and is further subdivided into forms containing sialic acid (negatively charged) and those without sialic acid (neutral).

In the present study, I examined the structure of oligosaccharide antennae located on the surface of apoB-100 and found that they are composed predominantly of N-linked oligosaccharides, i.e., two forms of complex type and five forms of high-mannose type. No O-l inked oligosac-charide was found.^15^ The proposed structures should be confirmed from several points of view such as the number of antennae, the number of sialic acids, etc. The types of antennae determined were as described in the Result section (Figs. 2, 3).It should be noted that neither tri-antennary nor tetra-antennary oligosaccharides are located on apoB-100.In bi-antennary oligosaccharides, the extent of sialylation could be determined as described without any further treatments. Since the DP shift by treatment with HEXase was small, there was no possibility that polylactosamine, i.e., a repeating of galactose and an N-acetylhexosamine unit, were attached to the antennae. Also, since the oligosaccharides were completely digested by HEXase, there was no possibility of the presence of a bisecting N-acetylhexosamine. The electrophoretic pattern also showed that there was no possibility of the presence of β (1-3) galactose instead of β(1-4) galactose. I did not perform any further analyses on the core structure, including fucose linkage as a Lewis structure. Because of relatively low concentrations of other types of antennae, if any, the endoglycosidase digestions did not result in detectable recovery by electrophoresis using an OLIGO Profiling Gel. Taniguchi et al. reported that the sugar chains consist of high-mannose-type oligo-saccharides, monosialylated and disialylated bi-antennary complex-type oligosaccharides, and small amounts of hybrid-type oligosaccharides.^2^ Garner et al. reported that 15 peaks were resolved by normal phase high-performance liquid chromatography and matrix-assisted laser desorption/ ionization time of flight mass spectrometry.^16^ However, there is a possibility that some of the elements were artificially produced by the hydrazinolytic procedure.

Although previous studies suggest that the oligosaccharides may play a role in the regulation of degradation very early in the secretion pathway, their post-hepatic roles in the circulation still remain unknown. Most studies in the area have focused on the relationship between lipoprotein-associated sialic acid and the development of atherosclerosis, and the idea that modifications of the oligosaccharide could induce atherogenic properties on LDL remains controversial.^17–19^ Mel’ nichenko et al. showed that desialylation of various apoB-containing lipoproteins can occur in the circulation, that this process increasesthe ability of these particlesto stimulate accumulation of cholesterol in human aortic intima cells, i e., increasestheir atherogenic potential.^20^ Several studies supported the idea that desialylation may contribute to the premature development of atherosclerosis.^21–24^ On the other hand, Chappey et al. observed no differences in the sialic acid content of LDL in subjects with coronary stenosis as compared to unaffected control subjects,^25^ as observed by other groups.^26^ They also have described that desialylation may result from an artificial modification of LDL related to in vitro peroxidation. One of the significant findings in our study was that equal molecules of mono sialylated and disialylated bi antennary complex- type oligosaccharides are located on the surface part of apoB-100 (lane 2 of Fig. 1).The digestion of sugar chains by exoglycosidases and endogly co-sidases did not result in any changes in the susceptibility of LDL to oxidation. This result showed that the sugar chains did not play any significant roles, at least not in the oxidative processing of LDL. Also, LDL without monosaccharides such as sialic acid, galactose, and N- acetylglucosamine did not induce any significant effect on the expression of ICAM-1, VCAM-1, or ELMA-1.This is the first study to demonstrate the functional roles of the total and partial structures of oligosaccharides on the surface of apoB-100.

## References

[1] Sharon N. Nomenclature of glycoproteins, glycopeptides and peptidoglycans. Pure & ApplChem 1988;60:1389–94.

[2] Taniguchi T, Ishikawa Y, Tsunemitsu M, Fukuzaki H. The structures of the asparagine-linked sugar chains of human apolipoprotein B-100. Arch Biochem Biophys 1989;273:197–205.

[3] Berliner JA, Heinecke JW. The role of oxidized lipopreoteins in atherosclerosis. Free Radic Biol Med 1996;20:707–27.

[4] Steinberg D. Low density lipoprotein oxidation and its pathological significance. J Biol chem 1997:272:20963–6.

[5] Kalant N, McCormick S, Parniak MA. Effects of copper and histidine on oxidative modification of low density lipoprotein and its subsequent binding to collagen. Arterioscler Thromb 1991;11:1322–9.

[6] Ronald A, Patterson RA, Leake DS. Measurement of copper-binding sites on low density lipoprotein. Arterioscler Thromb Vasc Biol 2001;21:594–602.

[7] Lamb DJ, Mitchinson MJ, Leake DS. Transition metal ions within human atherosclerotic lesions can catalyse the oxidation of low density lipoprotein by macrophages. FEBS left 1995;374:12–6.

[8] Dustin ML, Springer TA Lymphocyte function-associated antigen-1 (LFA-1) interaction with intercellular adhesion molecule-1 (ICAM-1) is one of at least three mechanisms for lymphocyte adhesion to cultured endothelial cells. J cell Biol 1988:107:321–31.

[9] Wellicome SM, Thornhill MH, Pitzalis C, et al. A monoclonal antibody that detects a novel antigen on endothelial cells that is induced by tumor necrosis factor, IL-1, or lipopolysac-chartide. J Immunol 1990:144:2558–65.

[10] Thornhill MH, Haskard DO. IL-4 regulates endothelial cell activation by IL-1, tumor necrosis factor, or IFN-γ. J Immunol. 1990;145:865–72.

[11] Okada M,Sugita O, Miida T, Maatsuto T. Inano K. Effects of modified low density lipoprotein and hypoxia on the expression of endothelial lwukocyte adhesion molecule-1. Presse Med 1995;24:483–8.

[12] Okada M, Matsuto T, Miida T, Inano K. Differences in the effects of cytokines on the expression of adhesion molecules in endothelial cells. Ann Med Interne 1997;148:125–9.

[13] Bruni S, Martinesi M, Stio M, et al. Effects of surface treatment of ti-6A1-4V titanium alloy on biocompatibility in cultured human umbilical vein endothelial cells. Acta Biomater 2005;1:223–34.

[14] Fukuta K, Asanagi M, Makino T. Structural control of sugar chains in animal cells. Trend Glycosci Glycotechnol 2001;13:395–405.

[15] Luo B, Soesanto Y, McClain DA. Protein modification by O-linked GlcNAc reduces angiogenesis by inhibiting Akt activity in endothelial cells. Arterioscler Thromb Vasc Biol 2008;28:651–7.

[16] Garner B, Harvey DJ, Royle L, et al. Characterization of human apolipoprotein B100 oligosaccharides in LDL subfractions derived from normal and hyperlipidemic plasma: deficiency of α-N-acetylneuraminyl-lactosyl-ceramide in light and small dense LDL particles. Glycobiology 2001;11:791–802.

[17] Bartlett AL, Stanley KK. All low density lipoprotein particles and partially desialylated in plasma. Atherosclerosis 1998;138:237–45.

[18] Millar JS. The sialylation of plasma lipoproteins. Atherosclerosis 2001;154:1–13.

[19] Millar JS, Anber V, Shephered J, Packard CJ Sialic acid-containing components of lipoproteins influence lipoprotein-proteoglycan interactions. Atherosclerosis 1999;145:253–60.

[20] Mel’nichenko AA, Tertov VV, Ivanova OA et al. Desialylation decreases the resistance of apoB-containing lipoproteins to aggregation and increases their atherogenic potential. Bull Exp Bio Med 2005;140:51–4.

[21] Harada LM, Carvalho MDT, Passarelli M,. Quintão ECR. Lipoprotein desialylation simultaneously the cell cholesterol uptake and impairs the reverse cholesterol transport system: in vitro evidences utilizing neuraminidase-treated lipoproteins and mouse peritoneal macrophages. Atherosclerosis 1998;139:65–75.

[22] Lindbohm N, Gylling H, Miettinen TE, Miettinen TA, Statin treatment increases the sialic acid content of LDL in hypercholesterolemic patients. Atherosclerosis 2000;151:545–50.

[23] Bartlett Al, Greqal T, Angelis ED, Myers S, Stanley KK, Role of the macrophage galactose leectin in the uptake of desialylated LDL. Atherosclerosis 2000;163:219–30.

[24] Terto VV, Kaplun VV, Sobenin IA, Orekhov AN. Low-density lipoprotein modification occurring in human plasma possible mechanism of in vivo lipoprotein desialylation as a primary step of atherogenic modification. Atheroscerosis 1998;138:183–95.

[25] Chapey B, Beyssen B, Foos E, et al. Sialic acid content of LDL in coronary artery disease: no evidence of desialylation in subjects with coronary stenosis and increased levels in subjects with extensice atherosclerosis and acute myocardial infraction, relation between desialylation and in vitro peroxidation. Artherioscler Thromb Vasc Biol 1998;18:876–83.

[26] Melaärvi N, Gylling H, Miettinen TA, Sialic acids and the metabolism of low density lipoprotein. J Lipid Res 1996;37:1625–31.

