## Supplemental Figures for "Sugar chain structure of apolipoprotein B-100 and its role in oxidation"

**Supplementary Table 1.** Sequences of Primers used in Real-Time RT-PCR

| Gene | Accession Number | Sequence |
| --- | --- | --- |
| ICAM-1 | NM_000201.2 | |
| | Forward primer | 5′-AACTGACACCTTTGTTAGCCACCTC-3′ |
| | Reverse primer | 5′-CCCAGTGAAATGCAAACAGGAC-3′ |
| VCAM-1 | NM_001078.2 | |
| | Forward primer | 5′-CGAAAGGCCCAGTTGAAGGA-3′ |
| | Reverse primer | 5′-GAGCACGAGAAGCTCAGGAGAAA-3′ |
| ELAM-1 | NM_000450.2 | |
| | Forward primer | 5′-ATGCCTGTGTGAGCAAGCATTTA-3′ |
| | Reverse primer | 5′-AGGCTAGAGCAGCTTTGGCAATTA-3′ |
| β-actin | NM_001101.3 | |
| | Forward primer | 5′-TGGCACCCAGCACAATGAA-3′ |
| | Reverse primer | 5′-CTAAGTCATAGTCCGCCTAGAAGCA-3′ |

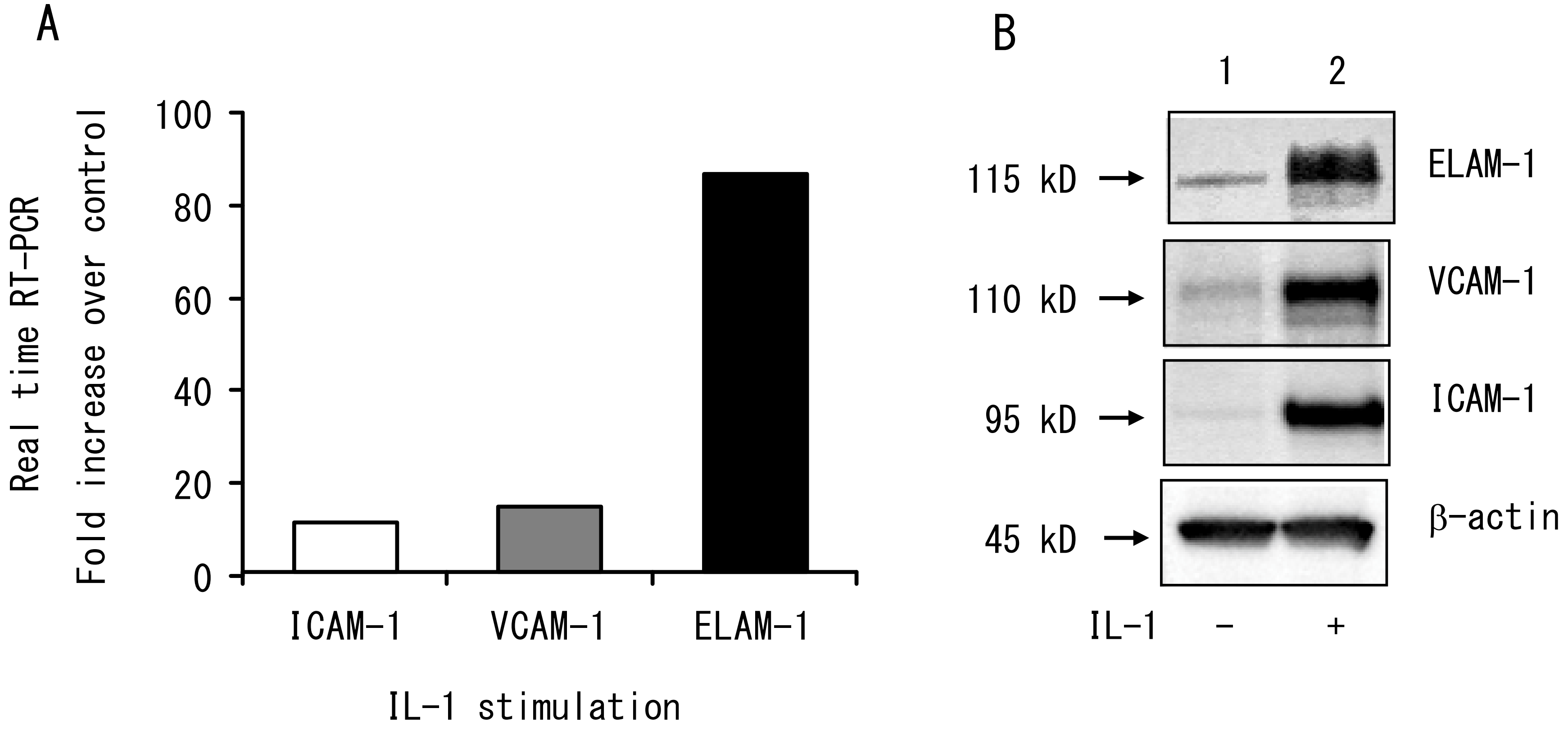


**Supplementary Fig. 1.** Effects of IL-1 stimulation on adhesion molecule expressions. Cultured coronary artery endothelial cells were simulated with 10 ng/mL IL-1α for 24 h in order to have positive controls for mRNA and protein analyses. ICAM-1, VCAM-1, and ELAM-1 were constitutively expressed on the cells and super-induced by the cytokine treatments. β-actin was used as internal control. **A**, Results of real time RT-PCR. **B**, Western blots.
